# Age dependent changes in circulating Tfh cells influence the development of functional antibodies to malaria in children

**DOI:** 10.1101/2021.12.12.472299

**Authors:** Jo-Anne Chan, Jessica R Loughland, Lauren de la Parte, Satomi Okano, Isaac Ssewanyana, Mayimuna Nalubega, Felistas Nankya, Kenneth Musinguzi, John Rek, Emmanuel Arinaitwe, Peta Tipping, Peter Bourke, Dean Andrew, Nicholas Dooley, Arya SheelaNair, Bruce D Wines, Mark Hogarth, James Beeson, Bryan Greenhouse, Grant Dorsey, Moses Kamya, Gunter Hartel, Gabriela Minigo, Margaret Feeney, Prasanna Jagannathan, Michelle J Boyle

## Abstract

T-follicular helper (Tfh) cells are key drivers of antibodies that protect from malaria. However, little is known regarding the host and parasite factors that influence Tfh and functional antibody development. Here, we use samples from a large cross-sectional study of children residing in an area of high malaria transmission in Uganda to characterize Tfh cells and functional antibodies to multiple parasites stages. We identify a dramatic re-distribution of the Tfh cell compartment with age that is independent of malaria exposure, with Th2-Tfh cells predominating in early childhood, while Th1-Tfh cell gradually increase to adult levels over the first decade of life. Functional antibody acquisition is age-dependent and hierarchical acquired based on parasite stage, with merozoite responses followed by sporozoite and gametocyte antibodies. Antibodies were boosted in children with current infection, and were higher in females. The children with the very highest antibody levels had increased Tfh cell activation and proliferation, consistent with a key role of Tfh cells in antibody development. Together, these data reveal a complex relationship between the circulating Tfh compartment, antibody development and protection from malaria.

## INTRODUCTION

*Plasmodium falciparum* malaria remains a leading cause of morbidity and mortality in children, and malaria control programs have been further negatively impacted by the SARS-CoV-19 pandemic (Organisation”, 2020). No vaccine yet approaches the efficacy of naturally acquired immunity, which provides nearly complete protection against symptomatic disease in older children and adults living in highly endemic areas. In such settings, the response to infection evolves slowly with age from acute, high-density, symptomatic infections in young children, to infections that are mostly lower density and asymptomatic in older children and adults. Immune development independently increases with both cumulative exposure and host age (Rodriguez-Barraquer et al., 2016, 2018). A better understanding of the mechanisms underpinning immunity and the factors that influence how immunity is acquired is needed to inform the development of improved vaccines and therapeutics that target the immune response.

An important mediator of protective immunity are antibodies that target the parasite to limit replication and parasite burden. Antibody development is driven by CD4 T-follicular helper (Tfh) cells that play key roles in germinal centres to activate naïve B cells and induce memory and antibody-producing B cells (Crotty, 2019)Circulating Tfh cells that resemble germinal center Tfh have been identified from peripheral blood (Vella et al., 2019), and can be subdivided into Th1-, Th2- and Th17-like subsets based on CXCR3 and CCR6 chemokine expression (Locci et al., 2013; Morita et al., 2011). Different Tfh subsets have been associated with antibody induction, depending on the infecting pathogen or vaccine, and context of exposure. During experimental human malaria in previously malaria-naive adults, all subsets of Tfh cells are activated (Chan et al., 2020). However only activated Th2-Tfh cells, and not other Tfh subsets, were associated with the induction of anti-malarial antibodies (Chan et al., 2020). We have recently shown that activation of Tfh cells during malaria is influenced by age, with adults having greater Tfh activation during infection, particularly within the Th2-Tfh cell subset (Oyong et al., 2021). In contrast, in children with malaria, activation is restricted to Th1-Tfh cells (Obeng-Adjei et al., 2015; Oyong et al., 2021). The impact of this age-dependent activation of Tfh cells on antibody development in malaria is unknown.

While antibodies have been recognised as essential mediators of protection from human malaria for over 50 years (Cohen et al., 1961), more recent studies have shown that the functional capacity of antibodies can be a better correlate of protective immunity than their quantity/titre (Beeson et al., 2019). Antibodies mediate a variety of effector functions, including fixing complement proteins and interacting with Fcγ receptors (FcγR) to promote opsonic phagocytosis and antibody-dependent cellular cytotoxicity. The importance of these antibody effector mechanisms is established across all parasite life stages. For example, complement fixing antibodies targeting liver and blood stages have been associated with protection from malaria (Boyle et al., 2015; Kurtovic et al., 2018; Reiling et al., 2019), and are a critical mediator of transmission-blocking immunity (Healer et al., 1997; Read et al., 1994; Williamson et al., 1995). Opsonic phagocytosis and FcγR-dependent functions have also been associated with protection from liver and blood stage infection (Feng et al., 2018; Hill et al., 2013; Kurtovic et al., 2020a; Osier et al., 2014). Little is known regarding the relative kinetics of acquisition of specific functional antibodies across different stages of the parasite as immunity develops.

Due to the central role of Tfh cells in antibody development, targeting Tfh to maximize the induction of functional antibodies may be key to achieving highly efficacious malaria vaccines (Beeson et al., 2019; Kurtovic et al., 2020b). Thus, it is important to identify factors that influence Tfh cell development, particularly in children with high malaria exposure, and to determine how Tfh development and activation influence antibody acquisition. Here, we examined Tfh cells and functional antibodies against multiple parasites stages in Ugandan children ages 0-10 with high malaria exposure, and investigated the impact of age and malaria on immune development. Our findings reveal important host and malaria impacts on both Tfh cells and antibody responses that inform vaccine development.

## RESULTS

### Circulating Tfh cell subsets change dramatically with age, independent of malaria

We analysed Tfh cells in children enrolled in a longitudinal observational cohort in Nagongera, eastern Uganda (Kamya et al., 2015). For the present study, samples were collected between January and April 2013 during a period of high malaria burden, with an average entomological inoculation rate of 215 infectious bites per person per year (Kilama et al., 2014). Tfh cells were quantified by flow cytometry in 212 children aged 0-11 years (54% male) (**Supplementary Figure S1)**. Consistent with the very high exposure in this cohort, 42% of children had asymptomatic *P. falciparum* infection at the time of sampling. Individual *P. falciparum* infection risk was assessed by household mosquito exposure (mosquito counts performed in individuals’ homes, monthly), which correlates with infection in this cohort (Boyle et al., 2017; Kilama et al., 2014; Rodriguez-Barraquer et al., 2018)

Tfh cells (CXCR5+PD1+) increased as a proportion of all CD4 T cells with age, (**Figure 1A**). The proportion of Tfh cells was not impacted by current asymptomatic parasitemia, household mosquito exposure, nor sex (**Figure 1B-D**). Unexpectedly, within the Tfh cell compartment, there was a dramatic redistribution of Tfh subsets with age, with an increase in Th1-Tfh and a decrease of Th2-Tfh cells between ages 0-7 years (Tfh subsets analysed based on CXCR3 and CCR6 expression. Th1-CXCR3+CCR6-, Th2 – CXCR3-CCR6-, Th17 – CXCR3-CCR6+, **Figure 1E**). Current infection and sex had no impact on Tfh subset distribution (**Figure 1F/H**). There was a suggestion that household mosquito exposure was associated with the proportion of Th17-Tfh cells, however this was not dose dependent (reduced Th17-Tfh in children with >40-80 compared to those with >80 household mosquitos/day (Dunn test FDR adjusted p = 0.04, **Figure 1G**). FoxP3+ Tfh (TfReg cells), have important roles in regulating germinal centre Tfh cells(Clement et al., 2019). However, how these responses related to FoxP3+ Tfh cells in the periphery is unclear, and here the proportion of FoxP3+ Tfh cells was not modified by age, current infection, household mosquito exposure but there was trend to higher FoxP3+ Tfh cells in males (**Supplementary Figure S2**).

**Figure 1:**
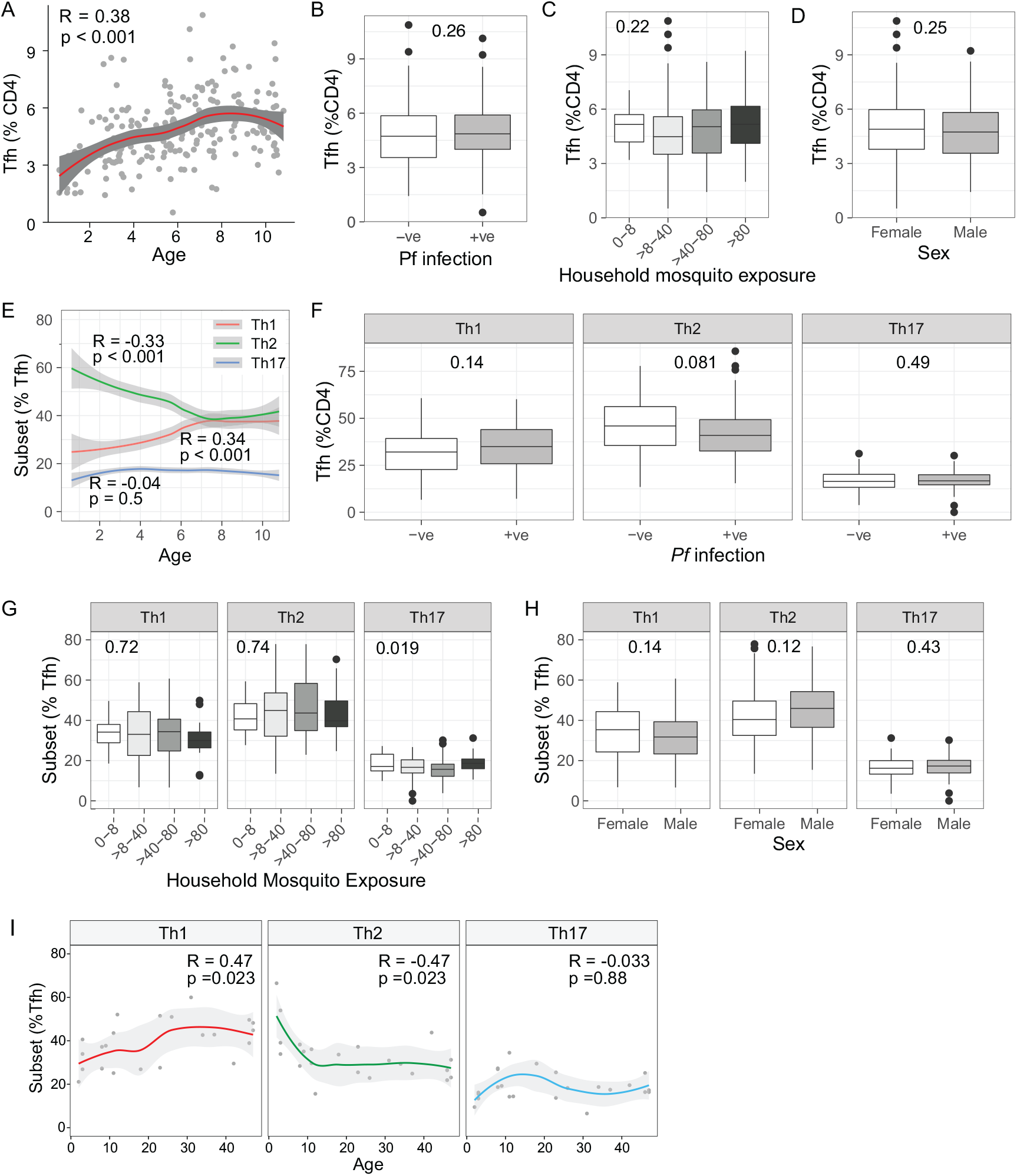
Impact of age and infection on Tfh cells subset composition. **A-H)** Tfh cells cells were analysed by flow cytometry in n = 212 Ugandan children. **(A-D**) Tfh cells were identified as CXCR5+PD1+ CD4 T cells. The relationship of the proportion of Tfh cells within the CD4 T cell compartment with age **(A)**; current asymptomatic P. falciparum infection **(B)**; household mosquito exposure **(C)**; and sex **(D)**. (**E-G**)Tfh cells were analysed based on CXCR3 and CCR6 expression into Th1 (CXCR3+CCR-), Th2 (CXCR3-CCR6-) and Th17 (CXCR3-CCR6+) subsets. The relationship of the proportion of Tfh subsets within the Tfh compartment with age **(E)**; current asymptomatic P. falciparum infection **(F)**; household mosquito exposure **(G)**; and sex **(H). I)** Tfh cells subsets were analysed by flow cytometry in malaria naïve children and adults (n=23) from a low infection burden setting in Australia and the relationship with age assessed. **A/E/I)** Solid lines are LOESS fit curves with 95% confidence interval. Spearman rho and p is indicated. **B/D/F/H)** Mann Whitney U test indicated. **C/G)** Kruskal Wallis indicated. See also Supplementary Figure S1 and S2.

To determine whether the change in Tfh subset distribution within our Ugandan cohort was driven by age or unaccounted for levels of malaria infection (which are co-linear with age in high transmission settings) and/or overall infectious diseases burden, we assessed Tfh cells within healthy, malaria naïve children and adults from low infectious diseases burden community within Australia. Consistent with an age driven change in the distribution of Tfh subsets, there was a marked decrease in Th2-Tfh cells in children, which then stabilised into adulthood. Concurrently, Th1-Tfh cells increased in children, and then stabilised into adulthood (**Figure 1I**). Taken together, these data suggest that the Tfh subset distribution undergoes marked changes during childhood, independent of malaria infection.

### Tfh cell activation and proliferation are influenced by both age and *P. falciparum* infection

During activation, Tfh cells upregulate inducible co-stimulator (ICOS) and the intracellular proliferation marker Ki67, which are required for Tfh cell function and development (Akiba et al., 2005; Herati et al., 2017; Rasheed et al., 2006). As such, we assessed the impacts of age and malaria on ICOS and Ki67 expression on Tfh cells. Both ICOS+ and Ki67+ expression on Tfh cells decreased with age (**Figure 2A/B**). Asymptomatic parasitemia was associated with a significant increase in Ki67 but not ICOS on Tfh cells (**Figure 2C/D**). High household mosquito exposure (>80 mosquitos/household/night) was associated with increased ICOS and Ki67 (Dunn test FDR adjusted ICOS >80 compared to 0-8 p=0.05, compared to >8-40 p=0.04, Ki67 >80 compared to 0-8 p=0.02; compared to >8-40 p=0.02, compared to >40-80 p=0.009, **Figure 2E/F**). Sex had no impact on ICOS or Ki67 expression (p=0.15 and p=0.72 respectively).

**Figure 2:**
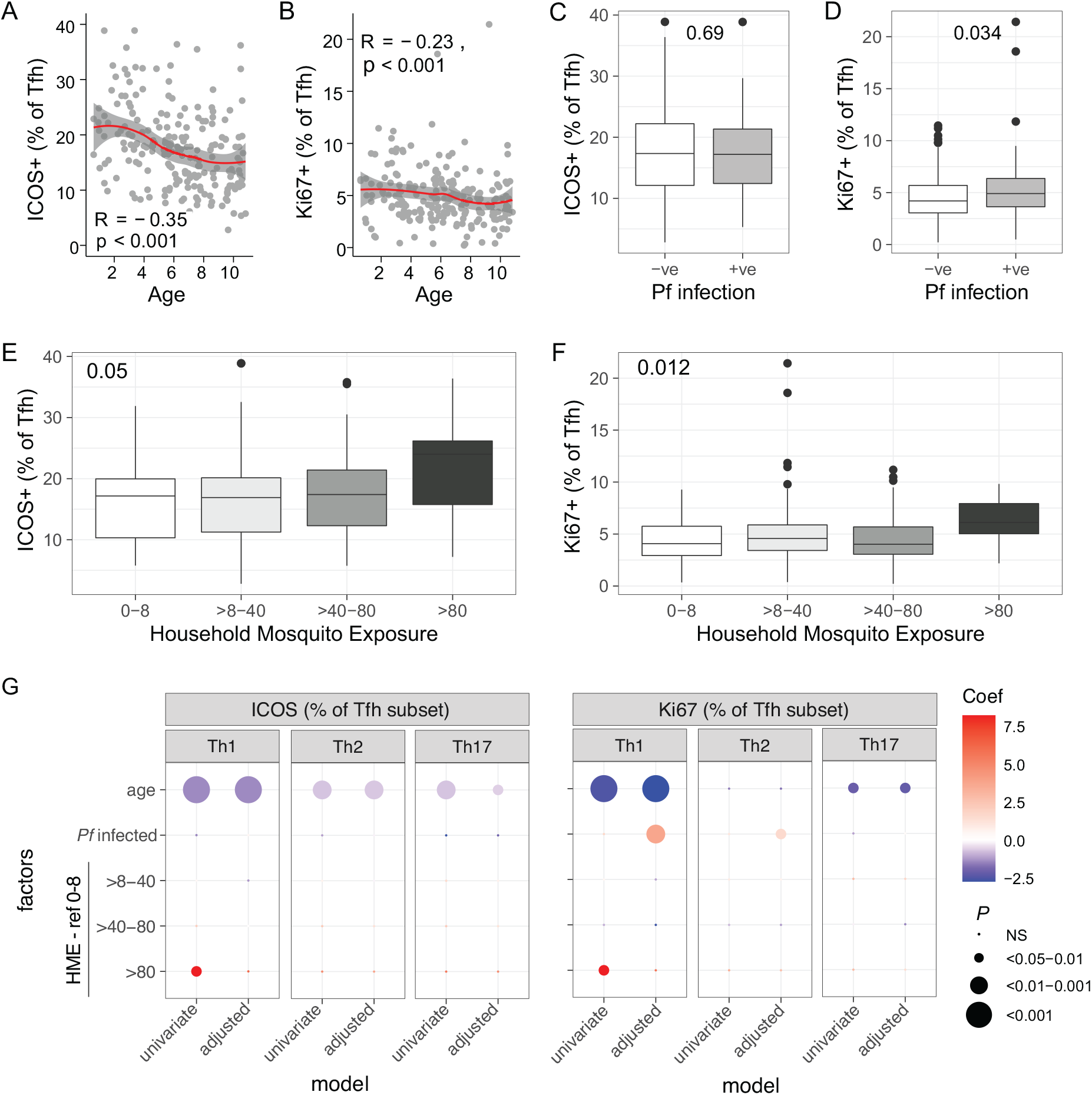
Impact of age and malaria on Tfh cell activation and proliferation. Activation and proliferation of Tfh cells was measured by ICOS and Ki67 expression in 212 Ugandan children. **A/B)** The relationship between ICOS+ and Ki67+ on Tfh cells with age. Line is LOESS Spearmans rho and p indicated. **C/D)** The relationship between ICOS and Ki67 and current asymptomatic infection Mann-Whitney U test indicated. **(E/F)** The relationship between ICOS and Ki67 on Tfh cells and household mosquito exposure. Kruskal Wallis indicated. **G)** Linear regression model coefficients for the relationship between ICOS andKi67 on each Tfh subset, and age, Pf infection and household mosquito exposure (HME).

Because of the interplay of age, current asymptomatic malaria infection and household mosquito exposure (Kilama et al., 2014), we used linear regression models to determine associations between these factors and ICOS/Ki67 expression in each Tfh subset. Age was associated with declining ICOS expression on Th1, Th2 and Th17 Tfh cells (**Figure 2G, Supplementary Table S1**). In contrast, Ki67 expression decreased with age on Th1- and Th17-, but not Th2-Tfh cells. Ki67 but not ICOS expression, was higher on both Th1- and Th2-Tfh cells in children with a current asymptomatic infection (**Figure 2G, Supplementary Table S1**). Sex had no impact on ICOS or Ki67 on any Tfh subset (data not shown). Thus, extending the previous observation that Tfh activation and proliferation during symptomatic malaria is restricted to Th1-like Tfh cells (Obeng-Adjei et al., 2015; Oyong et al., 2021), asymptomatic parasitemia is associated with increases in proliferation in both Th1- and Th2-Tfh cell subsets.

### Hierarchical acquisition of antibodies is dependent on parasite stage and antibody type and function

To investigate the relationship between Tfh cell compartment changes and antibody development, we comprehensively characterised malaria antibody responses in the cohort described above and an additional 50 children concurrently enrolled in the same study (total n=262). We measured antibodies to immunodominant blood stage (MSP2, AMA1), sporozoite stage (CSP) and gametocyte stage (Pfs230) antigens and evaluated the levels of IgG1, IgG3, and IgM along with functional antibodies that fixed complement, bound Fcγ receptors FcγRIIa and FcγRIII, and had the capacity to mediate opsonic phagocytosis by THP-1 monocytes (OPA). We have recently shown that opsonic phagocytosis by THP-1 monocytes is largely mediated by FcγRI (Feng et al., 2021). As expected, the prevalence and magnitude of both IgG and IgM antibodies increased with age (**Figure 3A/B**). Antibodies were first acquired to blood stage antigens MSP2 and AMA1, followed by sporozoite stage CSP, and finally gametocyte stage Pfs230. The large majority of children were seropositive to MSP2 and AMA1. In contrast, the majority of children had little antibody responses to gametocyte antigen Pfs230, with <50% seroprevalence for all antibody types even in older children (**Figure 3A**). For the blood stage antigen MSP2, IgG1 was the predominant subclass in young children, but switched to predominantly IgG3 with increasing age, consistent with previous publications (Dodoo et al., 2008; Duah et al., 2010; Noland et al., 2015; Taylor et al., 1998; Tongren et al., 2006). In contrast, IgG1 was the predominant subclass for PfAMA1 across all ages. For both blood stage antigens MSP2 and AMA1, opsonic phagocytosis antibodies were acquired first, followed by FcγRIII and then FcγRIIa-binding antibodies. C1q-fixing antibodies were less prevalent (25.8 and 33.3% seropositive for MSP2 and AMA1 respectively). Functional antibodies mediating OPA were also acquired first for CSP. However, antibodies targeting CSP were incapable of binding dimers of FcγRIIa or FcγRIII. There were very low levels of FcγRIIa and FcγRIII-binding, and no C1q-fixing antibodies targeting Pfs230.

**Figure 3:**
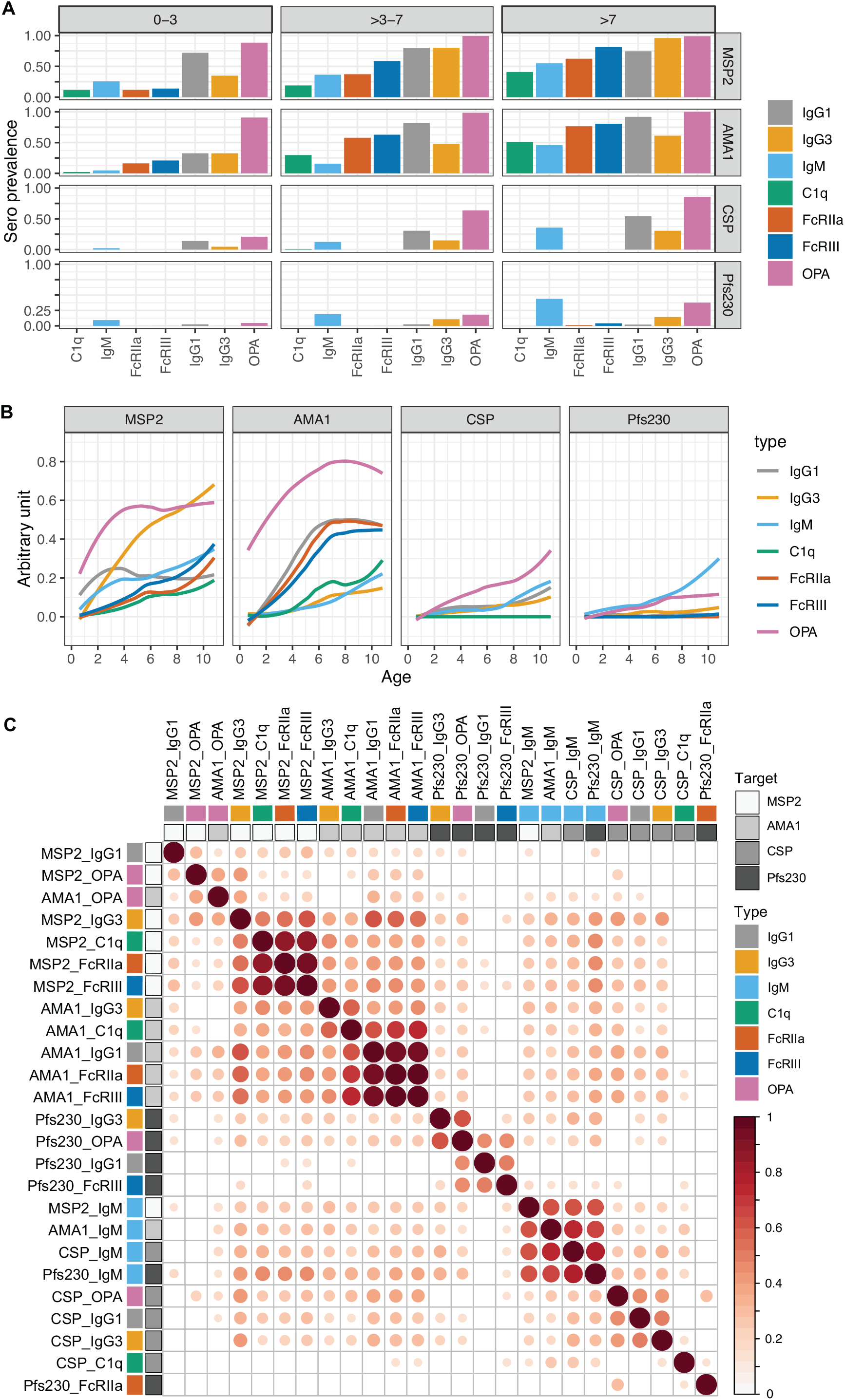
Seroprevalence and magnitude of antibodies with age. **(A)** Seroprevalence of antibody responses in children age 0-3, >3-7 and >7 years for IgG1, IgG3, IgM and functional antibodies that mediated complement fixation (C1q), cross linked FcRIIa and FcRIII and mediated opsonic phagocytosis (OPA) to blood stage (MSP2, AMA1), sporozoite stage (CSP) and gametocyte (Pfs230) antigens. Seroprevalence is defined as antibody levels greater than mean+3SD of negative controls from malaria naïve donors. **B)** Magnitude of antibodies to each antigen verses age with LOESS fit curves. Antibody magnitude is expressed as arbitrary units, which are calculated by thresholding data at positive seroprevalence levels and scaling data to a fraction of the highest responder. **C)** Pairwise Pearson’s correlations between the magnitudes of each antibody type/function to each antigen. Blank if correlation coefficient p>0.05. Rows and columns are clustered using hierarchical clustering with Ward.D method.

To investigate the relationship between antibody responses, we calculated the correlation between magnitudes of antibodies of each type to each antigen (**Figure 3C**). For MSP2, IgG3 levels were strongly correlated with C1q, FcRIIa and FcRIII, suggesting IgG3 may be a mediator of these functional responses. In contrast, for AMA1, IgG1 was more tightly correlated with C1q, FcRIIa and FcRIII than IgG3, suggesting that for AMA1 targeted antibodies IgG1 may be the dominant antibody driving functional responses. For CSP targeted antibodies, IgG3 and IgG1, and to a lesser extent IgM all correlated with opsonic phagocytosis levels, suggesting multiple antibody isotypes/subclasses may be involved with mediating phagocytosis of sporozoite stage parasites. In contrast, while IgM has recently been identified as a mediator of opsonic phagocytosis of merozoites (Hopp et al., 2021), MSP2 IgM levels were not correlated with MSP2 opsonic phagocytosis. Additionally, AMA1 IgM was only weakly correlated with opsonic phagocytosis, suggesting that different targets may be involved in IgM mediated opsonic phagocytosis of merozoites. IgM antibodies were tightly associated across all antigens (MSP2, AMA1, CSP and Pfs230), suggesting that IgM acquisition is con-current across all parasite live-stages. There was also a correlation between antibodies that mediate opsonic phagocytosis by monocytes to the blood-stage antigens MSP2 and AMA1. For antibodies targeting Pfs230, there was a strong correlation of IgG3 with opsonic phagocytosis and of IgG1 with FcRIII binding, suggesting different relationships between IgG subtypes and functions targeting this gametocyte antigen.

Taken together, these data highlight the complex relationships between antibody isotype/subtype and function, and the differential acquisition of antibodies across parasite stage and functional type. Our data suggest an age-driven hierarchical acquisition of antibodies, which is influenced by both parasite stage (merozoite>sporozoite>gametocyte) and function/subclass (for MSP2-OPA>IgG3/IgG1>FcRIII>FcRIIa> IgM>C1q; for AMA1-OPA>IgG1>FcRIII>FcRIIa>IgG3>IgM>C1q; for CSP – OPA>IgG1>IgM>IgG3>C1q; and for Pfs230 – IgM>OPA>IgG3>IgG1>FcRIIa>FcRIII).

### Antibody responses are higher in children with current infection and in females

Previous studies have consistently shown that antibody responses are boosted during infection. Consistent with this, the majority of antibody responses to blood stage antigens MSP2 and AMA1 were higher in children with a current asymptomatic *P. falciparum* infection (MSP2 IgG3, FcRIII, OPA; AMA1 IgG1, IgG3, C1q, FcRIIa, FcRIII, OPA, **Figure 4A**). There was also some boosting to non-blood stage antigens, with increased OPA antibodies to both CSP and Pfs230 during infection. IgM antibodies were not increased to any antigens in currently infected children, in contrast with previous findings by ourselves and others (Boyle et al., 2019; Walker et al., 2020). Household mosquito exposure was not associated with antibody responses (**Supplementary Figure S3**). Sex influenced multiple antibody responses, with higher IgG3 and C1q to MSP2 and AMA1, and FcRIIa to MSP2 in female children (**Figure 4B**).

**Figure 4:**
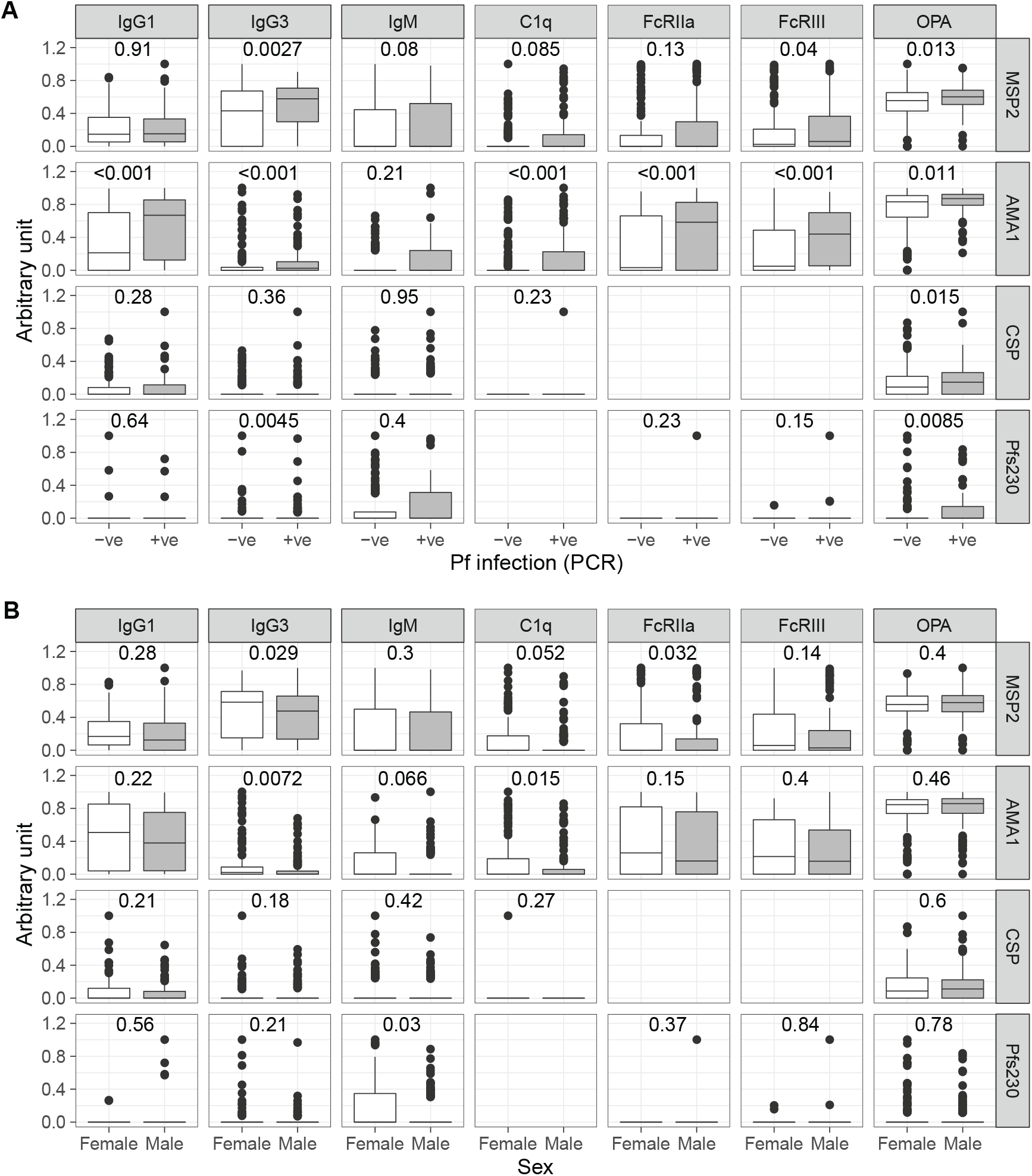
Relationships between antimalarial antibodies and current infection and sex. Magnitude of antibody responses of IgG1, IgG3, IgM and functional antibodies that mediated complement fixation (C1q), cross linked FcRIIa and FcRIII and mediated opsonic phagocytosis (OPA) to blood stage (MSP2, AMA1), sporozoite stage (CSP) and gametocyte (Pfs230) antigens. Antibody magnitude is expressed as arbitrary units, which are calculated by thresholding data at positive seroprevalence levels and scaling data to highest responder. **A)** Magnitude of antibodies in children with and without a current P. falciparum infected detected by PCR. **B)** Magnitude of antibodies in female and male children. Mann Whitney U tests indicated.

### Tfh activation and proliferation are associated with functional malaria-specific antibodies

To investigate whether Tfh phenotypes are associated with malaria-specific antibody development, we performed an unbiased principal components analysis (PCA) incorporating all Tfh and antibody responses from all individuals with complete data (n=211). This analysis identified four clusters of individuals (**Figure 5A**) that differed with respect to age and infection, but not household mosquito exposure and sex (**Figure 5B-E**). Children in cluster 1 were significantly younger than other clusters, followed by cluster 2 children, however there was no difference between clusters 3 and 4 (**Figure 5B**). Current infection was also associated with cluster (Chi-square test p=0.05), however infection between clusters 3 and 4 was comparable (**Figure 5C**). Proportion of Tfh cells (in CD4 T cells), the proportion of Tfh subsets (in Tfh cells), and ICOS and Ki67 expression (in total or Tfh subset) on clusters 1-3 was consistent with age driven changes, that was associated with increased Tfh cells within CD4 T cells, increased Th1-Tfh and decreased Th2-Tfh, and reduced ICOS and Ki67 (**Figure F-G**). In comparing cluster 3 to cluster 4 (which had comparable age distribution of individuals), cluster 4 had significantly reduced proportions of Th17-Tfh cells, increased ICOS on total Tfh and all subsets, and increased Ki67 on Th2-Tfh cells. Individuals in cluster 4 also had the very highest levels of cytophilic and functional antibodies compared to other clusters, including cluster 3. Together, PCA and clustering analysis is consistent with a strong impact of age on both Tfh and antibodies. However, this analysis also identifies two distinct clusters of older children, with individuals in cluster 4 with the highest antibodies also having increased ICOS expression on Tfh cells, and Ki67+ Th2-Tfh cells compared to similarly aged children in cluster 3. This finding is consistent with an important role of Th2-Tfh cell proliferation and Tfh activation in malaria antibody development (Bowyer et al., 2018; Chan et al., 2020; Minassian et al., 2021).

**Figure 5:**
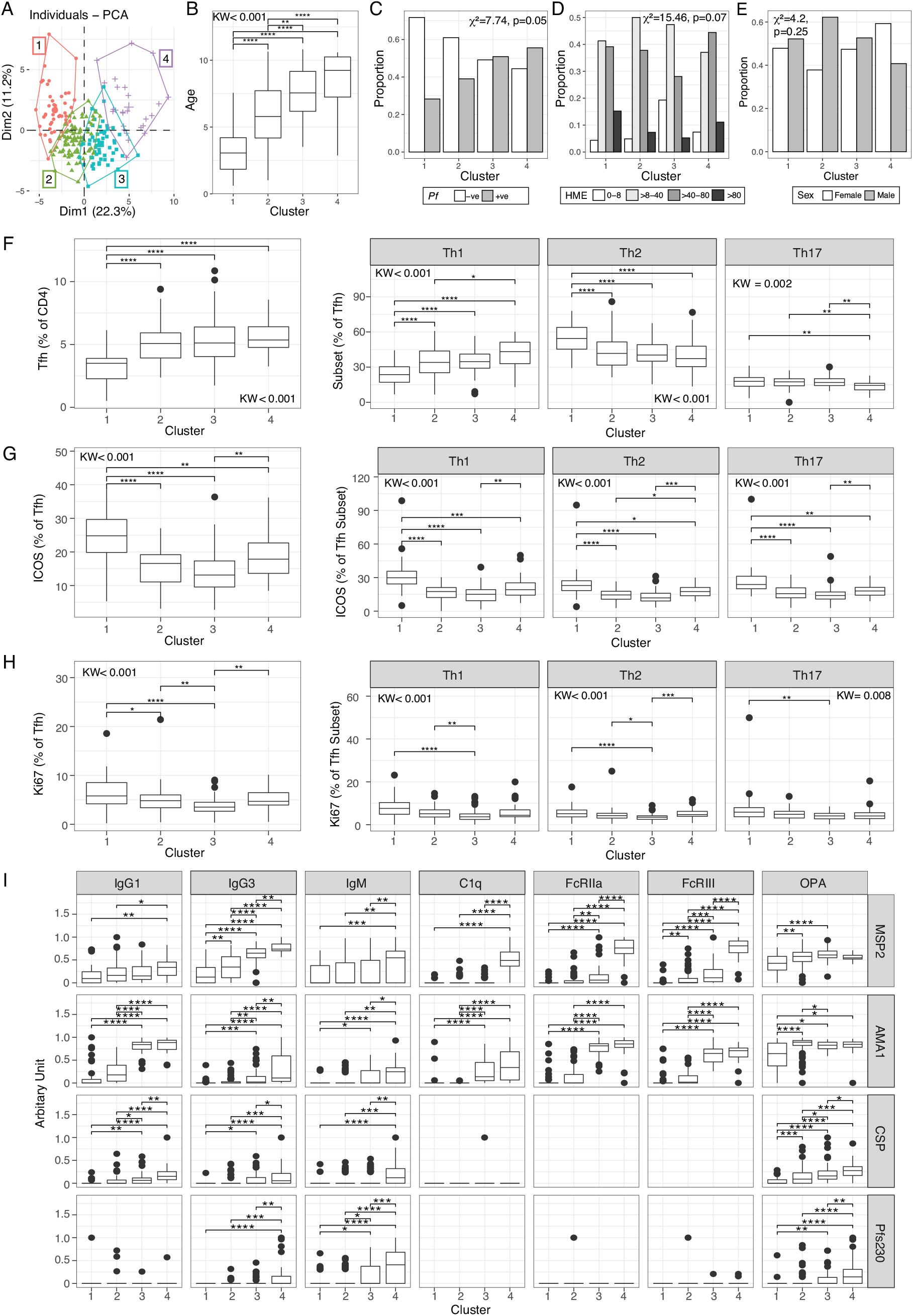
Principle component analysis and clustering of individuals. **A)** Tfh and antibody responses were analysed by PCA and individuals clustered by kmeans. **B)** Age, **C)** infection, **D)** household mosquito exposure (HME) and **E)** sex in kmeans clusters. **F)** Tfh cells and subsets in kmeans clusters. **G)** ICOS and **H)** Ki67 on total Tfh and Tfh cell subsets in kmean clusters. **I)** Antibody levels for each isotye/subclass and function in each kmean cluster. **B, F-I)** Kruskal Wallis and Dunn post analysis FDR adjusted indicated. **C-D)** Chi-square test indicated

### Associations of Tfh subsets and antibodies with protection from malaria

Finally, to determine the importance of Tfh cell subsets and antibodies in protective immunity to malaria, we measured their association with *P. falciparum* infection and probability of symptoms given infection in the following year. We first considered the odds of any *P. falciparum* infection (including both sub-microscopic and microscopic infection, and both asymptomatic and symptomatic infections). We found that higher proportions of Th17-Tfh cells, but no other Tfh subsets, were associated with reduced odds of any *P. falciparum* infection in the subsequent year of study. This association remained significant after controlling for age, current infection and household mosquito exposure (**Figure 6A, Supplementary Table S2**). For humoral responses, antibodies to blood stage antigens MSP2 and AMA1 were associated with an increased odds of infection in the following year. This is consistent with previous studies showing that antibodies to blood stage malaria parasites can accurately measure exposure risk (Greenhouse et al., 2011; Stanisic et al., 2015). Indeed IgG3, C1q, FcRIIa, FcRIII, OPA targeting MSP2 and IgG1, FcRIIa and FcRIII and OPA targeting AMA1 were all associated with increased odds of infection despite adjustment for age, current infection and household mosquito exposure (**Figure 6B, Supplementary Table S3**).

**Figure 6:**
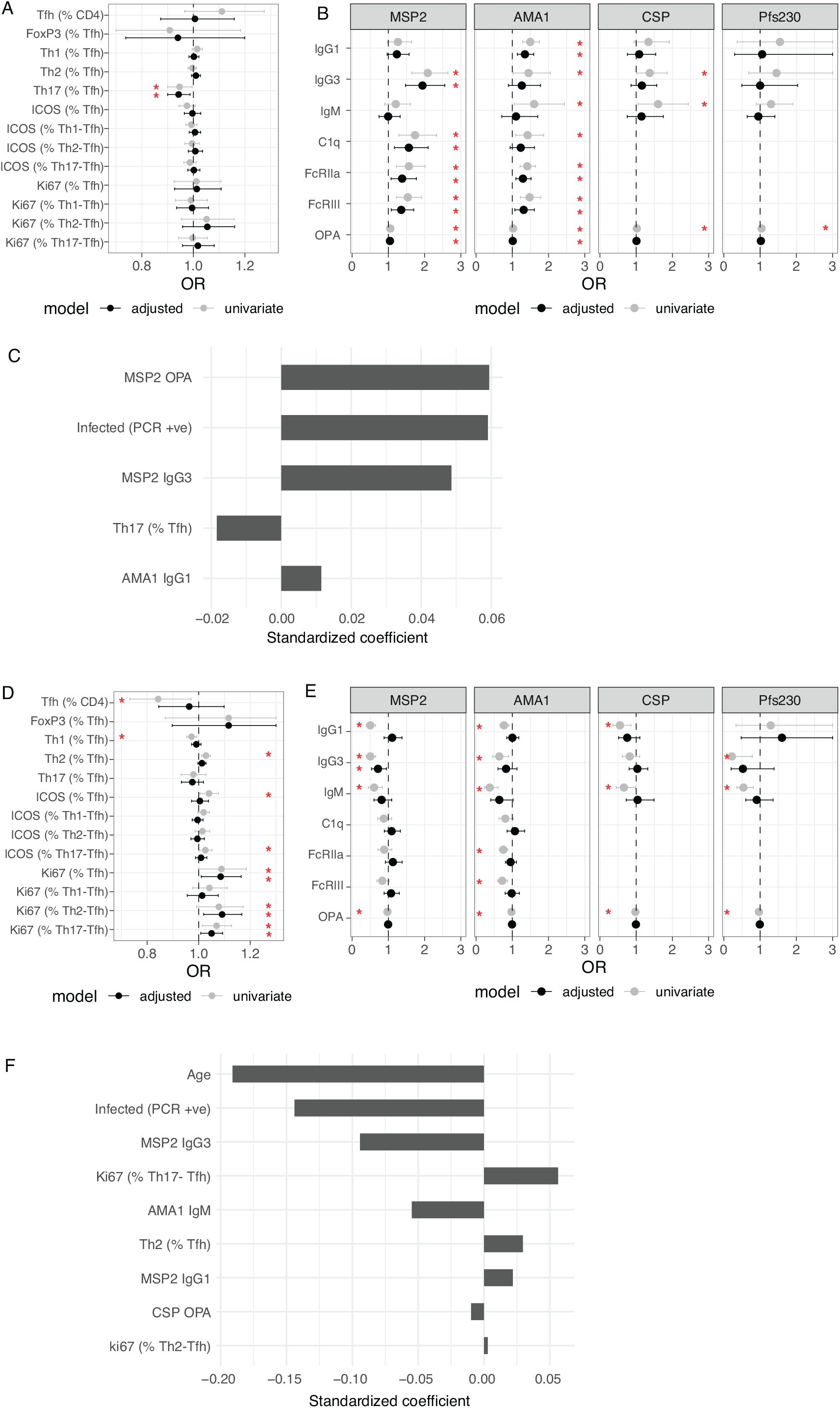
Association between Tfh cells, antibodies and malaria. Odds ratios for (**A**) Tfh subsets and **B)** antibodies with any infection (LAMP positive or blood smear positive) in the year following. **C)** LASSO Poisson regression standardized coefficients for responses selected as informative prognostic factors for incidence of any infection (LAMP positive or blood smear positive). Odd ratios for **D)** Tfh subsets and **E)** antibodies with symptomatic infection (blood smear positive with fever, given infection (LAMP positive or blood smear positive) in the year following. **F)** LASSO Poisson regression standardized coefficients for responses selected as informative prognostic factors for symptoms given infection. **A/B/D/F)** Unadjusted data and data adjusted for potential confounders are presented; in (**A/B**), data were adjusted for age (continuous), current infection detected by PCR and household mosquito exposure (HME), while in (**D/E**), data were only adjusted for age (continuous) and current infection. See also Supplementary Tables S2-5.

As an alternative approach, we modelled associations between both Tfh subsets and antibodies and the risk of infection using Least Absolute Shrinkage and Selection Operator (LASSO) regression modelling. This model is suited to selecting important components from multicollinear data and indicated that MSP2 OPA and IgG3, and AMA1 IgG1 antibodies, the proportion Th1-17-Tfh cells and current asymptomatic infection were important predictors that are associated with incidence of any density infection (**Figure 6C**).

We next assessed the association of Tfh subsets and antibodies with odds of symptomatic malaria when infected as a measure of protection from disease. Ki67 on total Tfh cells and Th2- and Th17-Tfh subsets (Ki67 % of Tfh, % of Th2-Tfh and %Th17-Tfh) were associated with increased odds of symptoms when infected after adjusting for age and current infection (**Figure 6D, Supplementary Table S4**). For antibody responses, MSP2 (IgG1, IgG3, IgM, OPA), AMA1 (IgG1, IgG3, IgM, FcRIIa, FcRIII, OPA), CSP (IgG1, IgG3, OPA) and Pfs230 (IgG3, IgM, OPA) were all associated with reduced odds of symptoms given infection in the year following. After adjusting for age and current infection, only MSP2 (IgG3) responses remained strongly associated with reduced odds of symptoms when infected (p=0.02, **Figure 6E, Supplementary Table S5**). Using LASSO modelling, MSP2 IgG3, AMA1 IgM and CSP OPA antibodies, along with age and asymptomatic infection were selected as factors that best predicted and were associated with reduced incidence of symptoms given infection. In contrast, Ki67 expression on Th17-Tfh and Th2-Tfh cells, and the proportion of Th2-Tfh cells were associated with increased incidence of symptoms given infection (**Figure 6F**). Together, these data reveal a complex relationship between the circulating Tfh compartment, antibody development, and protection from malaria. Our findings suggest an important role of cytophilic antibodies to blood stage antigens in protection from symptomatic malaria.

## DISCUSSION

Antibody induction during infection, including malaria, is driven by T-follicular helper CD4 T cells (Soon et al., 2021). However, little is known regarding the factors that influence Tfh cell development in children living in malaria-endemic settings, or their impact on the relative kinetics of acquisition of functional antibodies against different malaria parasite stages. Here, we show that the Tfh cell compartment changes dramatically with age, with a marked decrease in Th2-Tfh cells in the first 10 years of life in both malaria-exposed and malaria-naïve children. In malaria-exposed children, Tfh cell activation and proliferation also increased with age and were further influenced by current *P. falciparum*. In addition, we demonstrate that antibody acquisition with age followed a hierarchical order, and that age-dependent changes in antibody subclasses and functions differed with parasite stage and antigen target. Clustering analyses revealed that children with the highest levels of cytophilic antibodies had increased activation of Tfh cells and proliferating Th2-Tfh cells compared to similarly aged children. Importantly, cytophilic and functional antibodies to blood and sporozoite stage parasite proteins were associated with protection from symptoms among those with *P. falciparum* infection. Together, these results have implications for our understanding of Tfh cell biology in humans and antibody mediated immunity from malaria.

We found that in malaria exposed and naïve populations, the Tfh cell compartment undergoes marked changes with age, with a significant decline in Th2-Tfh and an increase in Th1-Tfh cells in children. To the best of our knowledge, this is the first report of this age dependent re-distribution of Tfh subsets in children. The mechanisms driving this dramatic age redistribution of Tfh subsets are unknown, but may be connected to the overall maturation of the immune response with age. Indeed, Th1 supporting cytokines and Th1 CD4 T cell responses to infection only reach levels of adult maturity around age 5-10 (Kollmann et al., 2012). While the specific mechanisms that mediate Th2-Tfh compared to Th1-Tfh development are largely unknown (Kim et al., 2018), this maturation of the immune system to support Th1 response may be important. How these changes to Tfh subsets impact immune development with age requires further investigation. Changes to Tfh subset distributions were not further impacted by current asymptomatic infection in our cohort. However, both Th1-Tfh and Th2-Tfh cells had increased proliferation in children with a current asymptomatic infection. In contrast, studies in Mali and Indonesia have shown that only Th1-Tfh cell subsets are activated among children with symptomatic malaria (Obeng-Adjei et al., 2015; Oyong et al., 2021). These differences may be due to underlying levels of inflammation between asymptomatic and symptomatic parasite infection, where IFN*γ* and TNF*α* are associated with skewing of Tfh to Th1-like subsets during *Plasmodium* infection (Obeng-Adjei et al., 2017; Ryg-Cornejo et al., 2016). In addition, multiple antibody responses were also boosted in asymptomatic parasitemia, consistent with Th2-Tfh cell activation and proliferation playing a role in antibody development (Chan et al., 2020). Further, clustering analysis revealed that children with the highest antibody levels had relatively higher activation of all Tfh subsets and proliferation of Th2-Tfh cells, compared to similarly aged children. This finding is consistent with an important role of Tfh cells, particularly Th2-Tfh cells, in antibody development in malaria (Chan et al., 2020).

Understanding the acquisition of antibodies that protect against symptomatic malaria is crucial to inform the development of efficacious vaccines. Our study, for the first time, measured the magnitude and function of antibodies against immunodominant antigens from different parasite life cycle stages (blood, sporozoites and gametocyte antigens). We found that antibodies in this cohort were first acquired to blood stage antigens such as MSP2 and AMA1, while antibodies to sporozoite CSP and gametocyte Pfs230 were acquired later. These findings are consistent with previous results in which we have shown functional antibodies that bind FcγRIIa and FcγRIII are significantly higher to merozoites compared to CSP antigen in Kenya populations (Feng et al., 2021). Differences in antibody acquisition across parasite stages are likely driven by antigen burden. Blood stages have the highest parasite burden due to ongoing replication of the parasite within red blood cells, and both MSP2 and AMA1 are abundantly expressed on the surface of merozoites (Gilson et al., 2006). In contrast, sporozoites are hidden from the immune system due to their gliding motility through hepatocytes. Similarly, gametocytes reside within red blood cells and emerge only within the mosquito midgut, thus reducing their capacity to stimulate an immune response. The extremely slow and infrequent acquisition of functional antibodies to sporozoite stages is consistent with the minimal levels of sterile protection against hepatocyte infection in naturally exposed populations (Tran et al., 2013). Similarly, low levels of functional antibodies targeting gametocyte stages suggest that transmission blocking immunity is low in children, and that children could remain a large reservoir of transmission despite their gradual development of immune protection from disease (Andolina et al., 2021). Alongside parasite stage, it is clear that the fine specificities at the subsets and functional levels are also influenced by specific characteristics of the antigen. Here, MSP2 IgG3 responses dominated and correlated with functional capacity of antibodies, in contrast to AMA1 where IgG1 antibodies are dominant. This finding is consistent with previous reports of IgG3 dominance to some but not all merozoite antigens and specific protein epitopes (Cavanagh et al., 2001; Dobaño et al., 2019; Duah et al., 2010; Richards et al., 2010). While the mechanisms mediating IgG3 or IgG1 dominance to specific parasite proteins is unknown, the presences of specific T-cell epitopes that induce strong IL10 responses alongside IFNγ has been implicated from experimental models (Tongren et al., 2005). Consistent with an important role of IL10 in skewing antibodies to IgG3, there is an age associated switch from IgG1 to IgG3 dominance for MSP2 antibodies seen here and in other studies (Duah et al., 2010; Taylor et al., 1998; Tongren et al., 2006), that coincides to the development of IL10/IFNγ co-producing malaria specific T cells within these same Ugandan children (Boyle et al., 2017)

In our study, multiple cytophilic and functional antibodies to blood and sporozoite stage antigens were associated with protection from symptomatic infection, consistent with previous studies (Boyle et al., 2015; Hill et al., 2016; Kurtovic et al., 2020b, 2020a; Osier et al., 2014; Reiling et al., 2019). Sex was also associated with antibody levels, with increased antibodies in female children. While sex-based differences in malaria antibodies have been understudied, this finding is consistent with a recent study in a different cohort of Ugandan children aged 7-16 where female children/adolescents had higher MSP1 and MSP2 antibodies compared to males (Apio et al., 2020). Indeed, in other infection and vaccination settings, females generally have greater antibody responses and global IgG levels and B cells (Fink and Klein, 2018; Klein and Flanagan, 2016). A recent study showed that female children have faster rates of parasite clearance compared to males (Briggs et al., 2020). Hence, sex-based differences in antibody responses may play an important role in protection from malaria.

There were several important limitations to our findings. First, samples were obtained from Ugandan children at a single cross-sectional timepoint. Although this study represents one of the largest studies to date, future studies will need to perform repeated assessments within children over time to determine the impact of age and malaria exposure on both antibody responses and T cell maturation. Second, the analysis of circulating Tfh responses evaluated bulk, and not antigen-specific Tfh cell responses (Oyong et al., 2021). Although we identified important associations between Tfh populations, age, antibody acquisition and protection, future studies are planned to assess the antigen-specific and functional capacity of Tfh cell response in this population.

In conclusion, in a large cohort of children living in a highly malaria-endemic setting, we describe dramatic redistribution of the Tfh cell compartment from Th2-Tfh to Th1 dominance with age, and associations between Tfh activation, proliferation and concurrent *P. Falciparum* infection. Additionally, we have identified hierarchical acquisition of functional antibodies driven by parasite stage and antibody type/subclass and function, with cytophilic and functional antibodies significantly associated with protection from the symptoms of *P. falciparum* infection. Young children are the most at risk of malaria, and are thus the target of vaccine interventions. As such, these findings are of relevance for our understanding of development of immunity in this population, and to vaccine development for children in endemic areas.

## METHODS

### Study population

#### Ugandan children cohort

Samples were obtained from 262 children enrolled in a longitudinal study by the East African International Centres of Excellence in Malaria Research conducted in the high transmission areas of Uganda (Boyle et al., 2017; Jagannathan et al., 2017; Kamya et al., 2015). This cohort consists of 100 households within the rural Nagongera sub-county in the Tororo district, where malaria transmission is holoendemic with seasonal peaks from October to January and April to July. In each randomly selected household, one adult caregiver (>20 years) and all children (eligibility of those aged 6 months to 10 years) were enrolled in the study.Upon enrollment all study participants were given an insecticide treated bed net and followed for all medical care at a dedicated study clinic. Children who presented with a fever (tympanic temperature >38.0 °C) or history of fever in the previous 24 hours had blood obtained by finger prick for a thick smear. If the thick smear was positive for *Plasmodium* parasites, the patient was diagnosed with malaria regardless of parasite density, and treated with artemether-lumefantrine. Routine assessments were performed in the study clinic every three months, including blood smears and dry blood spots to detect for parasite infection. Negative blood smears obtained at routine assessments were tested for the presence of submicroscopic malaria parasites using loop-mediated isothermal amplification (LAMP). Blood was drawn from each participant at a single cross-sectional timepoint between January and April 2013.

#### Malaria naïve cohort

PBMCs were collected from a healthy malaria-naive cohort of children (n=13, median age 8 IQR [3-13], 38% female) and adults (n=14, median age 39.5 IQR [25-43], 43% female) from a clinic of hospital outpatients. Volunteers were assessed by an on-site immunologist, where they were confirmed immunologically healthy and malaria-naïve.

### Ethics statement

Ethics approval for the use of human samples was obtained from the Uganda National Council of Science and Technology (HS1019), the institutional review boards of the University of California (11-05995) and Makerere University (2011-167), and the Alfred Hospital (#328/17), QIMR-Berghofer (P3444 and P3445), Stanford University (IRB 41197) and Menzies School of Health Research Human Ethics Research Committees (2012-1766). Written informed consent was obtained from all adult study participants and parents or legal guardians of the children.

### Measuring antibodies to recombinant *P. falciparum* antigens by ELISA

Recombinant MSP2 FC27 was expressed in *E. coli* as previously described (Stanisic et al., 2009). PfAMA1 (Boyle et al., 2019), full-length PfCSP (Kurtovic et al., 2019) and Pfs230D1M (Chan et al., 2019) were expressed in the mammalian HEK293 cell expression system as previously described.

The level of antibodies to recombinant *P. falciparum* antigens and merozoites were measured by ELISA as previously described <u>(Stanisic et al., 2009)</u>. Antigens were coated at 1μg/ml in PBS and incubated overnight at 4°C. Merozoites were coated at 1×10^7 cells per ml and incubated for 2h at 37°C. Plates were blocked with 1% casein in PBS (Sigma-Aldrich) for recombinant antigens and with 10% skim milk in PBS for merozoites, for 2h at 37°C. Following that, human antibodies (tested in duplicate) were added. For IgG detection, plates were incubated with a goat anti-human IgG HRP-conjugate (1/1000; Thermo Fisher Scientific). For IgG subclasses and IgM detection, plates were incubated with an additional step of mouse anti-human IgG1, IgG2, IgG3, IgG4 or IgM (1/1000; Thermo Fisher Scientific) for 1h at room temperature, followed by detection with a goat anti-mouse IgG HRP (1/1000; Millipore) for 1h at room temperature. Colour detection was developed using TMB liquid substrate (Sigma-Aldrich), which was subsequently stopped using 1M sulphuric acid. Antibodies were diluted with 0.1% casein in PBS for recombinant antigens and with 5% skim milk for merozoites. Serum dilution used for MSP2 FC27 was 1/250 for IgG subclasses and IgM, 1/100 for C1q and FcγR. Serum dilution used for PfAMA1 was 1/250 for IgG subclasses and IgM, 1/100 for C1q and 1/125 for FcγR. Serum dilution used for PfCSP was 1/100 for IgG subclasses, 1/250 for IgM and 1/100 for C1q. Serum dilution used for Pfs230D1M was 1/100 for all isotypes/subclasses and functions. PBS was used as a negative control and plates were washed thrice (with PBS with 0.05% Tween for recombinant antigens and with PBS only for merozoites) in between antibody incubation steps, using an automated plate washer (ELx405, BioTeck, USA). The level of antibody binding was measured as optical density at 450nm (for TMB) using the Multiskan Go plate reader (Thermo Fisher Scientific).

### Complement fixation assay

The capacity of human antibodies to fix complement C1q was measured as previously described (Boyle et al., 2015; Reiling et al., 2019). Briefly, antigen coating and blocking was performed as described above for standard ELISA. After incubation with human antibody samples, purified human C1q (10μg/ml; Millipore) was added as a source of complement for 30 min at room temperature, followed by a rabbit anti-C1q (1/2000; in-house) detection antibody and finally, a goat anti-rabbit IgG HRP (1/2500; Millipore) for 1h at room temperature. The level of C1q-fixation was developed using TMB liquid substrate (Sigma-Aldrich), reactivity was stopped using 1M sulphuric acid and measured as optical density at 450nm.

### Measuring antibodies that cross-link Fc receptors

Measuring antibodies that have the capacity to cross-link Fc receptors was performed as previously described (Feng et al., 2021; Kurtovic et al., 2020a; Wines et al., 2016). Briefly, antigen coating and blocking was performed as described above for standard ELISA. After incubation with human antibody samples, 50μL of biotinylated recombinant soluble dimers (0.2μg/ml of dimeric FcγRIIa, 0.1μg/ml of dimeric FcγRIII; expressed in-house using the HEK293 system) were added to the plates and incubated for 1h at at 37°C. Subsequently, a streptavidin HRP-conjugated antibody (1/10000; Thermo Fisher Scientific) was added for 1h at 37°C. Colour detection was developed using TMB liquid substrate and measured as optical density at 450nm.

### Opsonic phagocytosis of antigen-coated beads (OPA)

Measuring opsonic phagocytosis of antigen-coated beads was performed as previously described (Feng et al., 2021). Briefly, antigen-coated beads (5×10^7^ beads/ml) were incubated with serum samples for 1h at room temperature before washing and co-incubation with THP-1 monocytes for 20 min at 37°C. The proportion of THP-1 cells containing fluorescent-positive beads was acquired by flow cytometry (FACS Verse, BD Biosciences) and analysed using FlowJo software. Negative controls from non-exposed Melbourne residents were included in all assays.

### Flow cytometry

Ex vivo Tfh phenotype and activation was assessed by flow cytometry. PBMCs were thawed in 10% FBS/RPMI, and rested for 2 hours at 37 °C,5% CO_2_. In brief, 1M PBMCs were stained with surface antibodies to identify Tfh subsets, CD3, CD4, CXCR5 PD-1, CXCR3 and CCR6, activation markers included ICOS and CD38 (**Supplementary Table S7**). PBMCs were stained for 15 mins at RT, washed with 2% FBS/PBS, for intracellular markers, PBMCs were permeabilised with CytoFix/CytoPerm (BD) and 1 X Perm/Wash (BD) and stained with intracellular markers Ki67 and FoxP3. Samples were acquired on Aurora Cytek 3 laser instrument (Australian samples) or an Attune NXT Flow cytometer (Ugandan samples).

### *P. falciparum* culture

*P. falciparum* 3D7 isolates were maintained in continuous culture in RPMI-HEPES medium supplemented with hypoxanthine(370mM), gentamicin (30mg/ml), 25 mM sodium bicarbonate and 0.25% AlbuMAX II (GIBCO) or 5% heat-inactivated human sera in O+ RBCs from malaria-naive donors (Australian Red Cross blood bank). Cultures were incubated at 37°C in 1% O_2_, 5% CO_2_, 94% N_2_. Trophozoite and schizont stage parasites were purified by MACS separation (Miltenyl Biotec). Magnet purified infected RBCs (iRBCs) were stored at 1:2 in Glycerolyte 57 Solution (Baxter Healthcare Corporation) for in vitro stimulation assays.

### In vitro iRBC stimulation assay

Malaria-naïve PBMCs were stimulated at 1:3 (PBMC: iRBC) ratio for 5 days at 37 °C, 5% CO_2_. Cells were incubated in RPMI supplemented with 10% FBS. After 5 days, Tfh activation was assessed by flow cytometry.

### Statistical analyses

All statistical data analyses were performed using Prism 7 (GraphPad), STATA (version 15) and RStudio (R version 4.0.4). For analysis of antibody levels, antibody magnitude is expressed as arbitrary units, which are calculated by thresholding data at positive seroprevalence levels and scaling data to highest responder. Correlations between antibody variables and age were assessed using Pearson correlation for continuous variables. Clustering of correlations was performed with H-clustering with Ward.D method with R Hmisc package. To assess the association between Tfh cells and antibodies once controlling for age, lower limit of antibodies magnitudes were set at 0.001 and then log transformed before analysis in linear regression modelling. For clustering analysis of individuals, only those with complete Tfh and antibody data were included. All data was transformed into z-scores and then PCA performed in factoextra and FactoMineR. Kmeans cluster number chosen in ‘useful’ package and kmeans clusters identified with ‘stats; package in R.

To assess the odds of infection and disease in the year following with antibody and Tfh subsets, routine assessments with active case detection were performed in the study clinic every 3 months, including blood smears and dry blood spots to detect parasite infections. Negative blood smears obtained at routine assessments were tested for the presence of sub-microscopic malaria parasites using loop-mediated isothermal amplification (LAMP) (Hopkins et al., 2013). At the time of routine assessments, children were classified into four categories as described previously (Boyle et al., 2017; Jagannathan et al., 2017): (1) no evidence of parasite infection; (2) asymptomatic, submicroscopic (LAMP positive) infection; (3) asymptomatic, blood smear positive infection, or (4) symptomatic malaria defined as fever with a smear positive infection, with a window of 21 days prior to and 7 days following the routine visit to ensure capture of malaria episodes that were recently treated or infections that soon became symptomatic. To assess relationships between cross-sectional antibody responses and the prospective odds of infection measured at repeated, quarterly routine assessments, the odds of any infection (defined as either LAMP positive or blood smear positive with or without symptoms) were calculated using multilevel mixed-effects logistic regression, accounting for repeat measures within individuals and clustered on household to account for multiple children in each individual household. In multivariate analysis, odds ratios for infection risk were adjusted for age (linear), current asymptomatic infection and household mosquito exposure (categorical 0–8, >8–40, >40–80, >80 mosquitos/household/day). Household mosquito exposure was calculated as described previously (Boyle et al., 2017). For each individual, household mosquito exposure was calculated based on the mean household-level female Anopheles mosquito counts obtained from CDC light traps placed overnight (once per month) within the household of all trial participants (Kilama et al., 2014). Mean female Anopheles mosquito counts from the prospective 12 months were calculated used in analysis as a categorical variable (0–8, >8–40, >40–80, and >80 mosquitos/household/day), based on analyses of the relationship between household mosquito exposure and *P. falciparum* infection (Boyle et al., 2017). Similarly, to assess the odds of symptoms given infection (defined as cases if clinical symptoms when an infection was detected either by LAMP or blood smear) and odds of sub-patent infection given any infection (defined as LAMP positive only if LAMP or blood smear positive with or without symptoms) were calculated using multilevel mixed-effects logistic regression, accounting for repeat measures within individuals and clustered on household to account for multiple children in each individual household. In multivariate analyses, odds ratios were adjusted for age (continuous) and current infection.

LASSO Poisson regression modelling was used to select the most informative prognostic factors for 1) incidence of density infections and 2) incidence of symptoms when infected across the multiple visits during the study period. Variables assessed include age, gender, daily mosquito exposure rate (continuous in log and categorical 0-8,>8-40, >40-80, >80), and OD values from 27 antibodies: The optimum lambdas were selected using 5-fold cross validation that achieves the minimum Poisson deviance. Relative variable importance of selected variables was described using standardized coefficients. Only children with complete information were included in the analyses (complete case analyses, n=211). R statistical software version 4.0.2 with R package “glmnet” was used (Friedman et al., 2010).

## AUTHOR CONTRIBUTIONS

JAC, JRL, GM, PJ, MJB designed research study;

JAC, JRL, LdelP, ALP, AS, DA, ND conducted experiments;

JAC, JRL, SO, GH, MJB analysed data;

BDW, MH, JB provided essential reagents, technical expertise and contributed to assay development;

IS, MN, FN, KM, BG, PT, EA, PB, JR, MK, GD, MF, PJ conducted and supervised the clinical studies and sampling;

JAC, JRL, GM, PJ, MJB led manuscript preparation with feedback from all authors.

## ACKNOWLEDGEMENTS

We thank Australian Red Cross Blood Service for providing malaria naïve samples. We thank all staff of Infectious Disease Research Collaboration, Uganda. We thank all study volunteers and their guardians.

## FUNDING

This work was supported by the National Health and Medical Research Council of Australia (Early Career Fellowship 1125656, Career Development Award 1141278, Project Grant 1125656 and Ideas Grant 11819312 to MJB; Senior Research Fellowship 1077636 to JGB). Additional support was provided by NIH (U19 A108974 to GD, and U01 AI150741 to PJ). The Burnet Institute is supported by the National Health and Medical Research Council for Independent Research Institutes Infrastructure Support Scheme and the Victorian State Government Operational Infrastructure Support.

## Notes

### Competing Interest Statement

The authors have declared no competing interest.

