## Supplementary Figures and Tables for "Age dependent changes in circulating Tfh cells influence the development of functional antibodies to malaria in children"

### Supplementary Material

**Supplementary Table S1: Linear regression modelling outputs of the relationship of ICOS and Ki67 expression on Tfh subsets with age, *Pf* infection and household mosquito exposure**

| | | univariate | | | | adjusted * | | | |
|---|---|---|---|---|---|---|---|---|---|
| | **ICOS+** | **coef** | **95% CI** | | **p** | **coef** | **95% CI** | | **p** |
| Th1 | age | -1.367 | -1.872 | -0.861 | **<0.001** | -1.373 | -1.900 | -0.845 | **<0.001** |
| | Pf infection | -1.432 | -4.440 | 1.576 | 0.349 | 0.497 | -2.433 | 3.426 | 0.739 |
| | HME 0-8 (ref) | | | | | | | | |
| | >8-40 | 0.456 | -4.893 | 5.806 | 0.867 | -1.229 | -6.316 | 3.858 | 0.634 |
| | >40-80 | 2.119 | -3.343 | 7.582 | 0.445 | 0.123 | -5.085 | 5.331 | 0.963 |
| | >80 | 8.191 | 1.273 | 15.109 | **0.021** | 6.108 | -0.474 | 12.690 | 0.069 |
| Th2 | age | -0.679 | -1.089 | -0.269 | **0.001** | -0.644 | -1.076 | -0.212 | **0.004** |
| | Pf infection | -1.116 | -3.461 | 1.228 | 0.349 | -0.187 | -2.585 | 2.211 | 0.878 |
| | HME 0-8 (ref) | | | | | | | | |
| | >8-40 | 0.658 | -3.550 | 4.866 | 0.758 | -0.159 | -4.324 | 4.006 | 0.940 |
| | >40-80 | 2.205 | -2.091 | 6.502 | 0.313 | 1.233 | -3.031 | 5.497 | 0.569 |
| | >80 | 4.714 | -0.728 | 10.156 | 0.089 | 3.759 | -1.630 | 9.148 | 0.171 |
| Th17 | age | -0.680 | -1.137 | -0.222 | **0.004** | -0.553 | -1.034 | -0.073 | **0.024** |
| | Pf infection | -2.719 | -5.307 | -0.132 | 0.349 | -1.981 | -4.651 | 0.689 | 0.145 |
| | HME 0-8 (ref) | | | | | | | | |
| | >8-40 | 1.548 | -3.134 | 6.231 | 0.515 | 0.705 | -3.928 | 5.339 | 0.764 |
| | >40-80 | 2.217 | -2.560 | 6.994 | 0.361 | 1.229 | -3.510 | 5.967 | 0.610 |
| | >80 | 5.462 | -0.588 | 11.512 | 0.077 | 4.737 | -1.251 | 10.725 | 0.120 |
| | **Ki67+** | **coef** | **95% CI** | | **p** | **coef** | **95% CI** | | **p** |
| Th1 | age | -0.362 | -0.542 | -0.182 | **<0.001** | -0.436 | -0.620 | -0.251 | **<0.001** |
| | Pf infection | 0.883 | -0.155 | 1.920 | 0.095 | 1.475 | 0.452 | 2.499 | **0.005** |
| | HME 0-8 (ref) | | | | | | | | |
| | >8-40 | 0.510 | -1.345 | 2.365 | 0.589 | 0.055 | -1.722 | 1.832 | 0.951 |
| | >40-80 | 0.092 | -1.802 | 1.986 | 0.923 | -0.431 | -2.250 | 1.389 | 0.641 |
| | >80 | 2.790 | 0.391 | 5.189 | **0.023** | 2.059 | -0.240 | 4.358 | 0.079 |
| Th2 | age | -0.080 | -0.212 | 0.052 | 0.234 | -0.119 | -0.256 | 0.019 | 0.091 |
| | Pf infection | 0.686 | -0.051 | 1.422 | 0.095 | 0.838 | 0.073 | 1.603 | **0.032** |
| | HME 0-8 (ref) | | | | | | | | |
| | >8-40 | 0.652 | -0.678 | 1.983 | 0.335 | 0.555 | -0.773 | 1.884 | 0.411 |
| | >40-80 | 0.134 | -1.225 | 1.492 | 0.846 | 0.028 | -1.332 | 1.388 | 0.968 |
| | >80 | 1.379 | -0.342 | 3.100 | 0.116 | 1.157 | -0.562 | 2.876 | 0.186 |
| Th17 | age | -0.248 | -0.457 | -0.040 | **0.020** | -0.268 | -0.488 | -0.048 | **0.017** |
| | Pf infection | 0.048 | -1.132 | 1.228 | 0.095 | 0.451 | -0.769 | 1.670 | 0.467 |
| | HME 0-8 (ref) | | | | | | | | |
| | >8-40 | 1.217 | -0.901 | 3.335 | 0.259 | 0.904 | -1.213 | 3.020 | 0.401 |
| | >40-80 | 0.308 | -1.853 | 2.469 | 0.779 | -0.052 | -2.217 | 2.113 | 0.962 |
| | >80 | 1.219 | -1.518 | 3.956 | 0.381 | 0.793 | -1.942 | 3.529 | 0.568 |

* There was no evidence for an interaction between age and infection on ICOS or Ki67 expression within the linear model.

### Supplementary Material

#### Supplementary Table S2: Relationship between Tfh subsets and odds of infection

| | Univariate | | | | Adjusted | | | |
|---|---|---|---|---|---|---|---|---|
| **Tfh subset** | **OR** | **p** | **95% CI** | | **aOR** | **p** | **95% CI** | |
| Tfh (% of CD4) | 1.11 | 0.139 | 0.967 | 1.275 | 1.006 | 0.938 | 0.873 | 1.158 |
| FoxP3 (% of Tfh) | 0.908 | 0.475 | 0.697 | 1.183 | 0.94 | 0.62 | 0.737 | 1.199 |
| Th1 (% of Tfh) | 1.015 | 0.14 | 0.995 | 1.035 | 1.002 | 0.844 | 0.983 | 1.021 |
| Th2 (% of Tfh) | 0.996 | 0.644 | 0.979 | 1.013 | 1.01 | 0.261 | 0.993 | 1.027 |
| Th17 (% of Tfh | 0.946 | **0.031** | 0.9 | 0.995 | 0.943 | **0.012** | 0.9 | 0.987 |
| ICOS (% of Tfh) | 0.975 | 0.137 | 0.944 | 1.008 | 0.996 | 0.808 | 0.965 | 1.028 |
| ICOS (% of Th1-Tfh) | 0.992 | 0.468 | 0.971 | 1.014 | 1.006 | 0.621 | 0.984 | 1.028 |
| ICOS (% of Th2-Tfh) | 0.994 | 0.697 | 0.967 | 1.023 | 1.008 | 0.579 | 0.981 | 1.036 |
| ICOS (% of Th17-Tfh) | 0.986 | 0.256 | 0.963 | 1.01 | 1.002 | 0.899 | 0.978 | 1.025 |
| Ki67 (% of Tfh) | 1.012 | 0.803 | 0.924 | 1.107 | 1.013 | 0.772 | 0.927 | 1.108 |
| Ki67 (% of Th1-Tfh) | 0.99 | 0.765 | 0.93 | 1.055 | 0.995 | 0.874 | 0.935 | 1.059 |
| Ki67 (% of Th2-Tfh) | 1.051 | 0.32 | 0.953 | 1.158 | 1.054 | 0.277 | 0.958 | 1.16 |

N=211 children. OR: odds ratio; aOR: adjusted OR.

* Adjusted for age as a continuous variable, current *P. falciparum* infection detected by PCR, and household mosquito exposure as a categorical variable, 0-8, >8-40, >40-80, >80 mosquitos/household/day

**Supplementary Material**

**Supplementary Table S3: Relationship between antibodies and odds of Infection**

| | | | **Univariate** | | | **Adjusted *** | | | |
|---|---|---|---|---|---|---|---|---|---|
| Target | **Type** | **OR** | **p** | **95% CI** | | **aOR** | **p** | **95% CI** | |
| MSP2 | IgG1 | 1.266 | 0.073 | 0.978 | 1.638 | 1.235 | 0.091 | 0.968 | 1.575 |
| | IgG3 | 2.087 | **<0.001** | 1.648 | 2.643 | 1.941 | **<0.001** | 1.472 | 2.558 |
| | IgM | 1.202 | 0.222 | 0.895 | 1.613 | 0.992 | 0.958 | 0.743 | 1.326 |
| | C1q | 1.736 | **<0.001** | 1.292 | 2.331 | 1.565 | **0.003** | 1.168 | 2.097 |
| | FcRIIa | 1.568 | **0.001** | 1.221 | 2.013 | 1.380 | **0.012** | 1.074 | 1.773 |
| | FcRIII | 1.532 | **<0.001** | 1.226 | 1.916 | 1.354 | **0.009** | 1.079 | 1.701 |
| | OPA | 1.057 | **<0.001** | 1.036 | 1.079 | 1.044 | **<0.001** | 1.023 | 1.065 |
| AMA1 | IgG1 | 1.498 | **<0.001** | 1.286 | 1.745 | 1.349 | **<0.001** | 1.143 | 1.592 |
| | IgG3 | 1.450 | **0.036** | 1.024 | 2.053 | 1.262 | 0.185 | 0.895 | 1.780 |
| | IgM | 1.605 | **0.027** | 1.055 | 2.441 | 1.101 | 0.666 | 0.712 | 1.703 |
| | C1q | 1.427 | **0.009** | 1.092 | 1.866 | 1.231 | 0.131 | 0.940 | 1.612 |
| | FcRIIa | 1.412 | **<0.001** | 1.216 | 1.640 | 1.294 | **0.002** | 1.101 | 1.519 |
| | FcRIII | 1.480 | **<0.001** | 1.225 | 1.788 | 1.313 | **0.009** | 1.071 | 1.609 |
| | OPA | 1.023 | **<0.001** | 1.012 | 1.035 | 1.013 | **0.037** | 1.001 | 1.025 |
| CSP | IgG1 | 1.338 | 0.110 | 0.936 | 1.914 | 1.080 | 0.671 | 0.758 | 1.539 |
| | IgG3 | 1.371 | **0.041** | 1.013 | 1.855 | 1.159 | 0.339 | 0.857 | 1.566 |
| | IgM | 1.605 | **0.020** | 1.057 | 2.438 | 1.148 | 0.522 | 0.753 | 1.749 |
| | OPA | 1.018 | **0.004** | 1.006 | 1.031 | 1.009 | 0.184 | 0.996 | 1.022 |
| Pfs230 | IgG1 | 1.547 | 0.552 | 0.368 | 6.504 | 1.056 | 0.933 | 0.296 | 3.761 |
| | IgG3 | 1.449 | 0.332 | 0.685 | 3.067 | 1.005 | 0.990 | 0.496 | 2.035 |
| | IgM | 1.304 | 0.171 | 0.892 | 1.906 | 0.951 | 0.803 | 0.643 | 1.408 |
| | OPA | 1.043 | **0.008** | 1.010 | 1.076 | 1.024 | 0.117 | 0.994 | 1.055 |

N=261 children. OR: odds ratio; aOR: adjusted OR.

* Adjusted for age as a continuous variable, and household mosquito exposure as a categorical variable, 0-8, >8-40, >40-80, >80 mosquitos/household/day.

### Supplementary Material

**Supplementary Table S4: Relationship between Tfh subsets and odds of symptoms given infection**

| | Univariate | | | | Adjusted | | | |
|---|---|---|---|---|---|---|---|---|
| **Tfh subset** | **OR** | **p** | **95% CI** | | **aOR** | **p** | **95% CI** | |
| Tfh (% of CD4) | 0.843 | **0.017** | 0.732 | 0.97 | 0.963 | 0.577 | 0.845 | 1.099 |
| FoxP3 (% of Tfh) | 1.116 | 0.388 | 0.87 | 1.432 | 1.116 | 0.324 | 0.897 | 1.39 |
| Th1 (% of Tfh) | 0.972 | **0.005** | 0.953 | 0.992 | 0.99 | 0.307 | 0.972 | 1.009 |
| Th2 (% of Tfh) | 1.028 | **0.001** | 1.011 | 1.045 | 1.012 | 0.124 | 0.997 | 1.029 |
| Th17 (% of Tfh | 0.979 | 0.401 | 0.932 | 1.029 | 0.975 | 0.251 | 0.933 | 1.018 |
| ICOS (% of Tfh) | 1.04 | **0.026** | 1.005 | 1.076 | 1.005 | 0.785 | 0.972 | 1.038 |
| ICOS (% of Th1-Tfh) | 1.019 | 0.091 | 0.997 | 1.041 | 0.996 | 0.684 | 0.975 | 1.017 |
| ICOS (% of Th2-Tfh) | **1.014** | 0.32 | 0.986 | 1.043 | 0.995 | 0.731 | 0.97 | 1.022 |
| ICOS (% of Th17-Tfh) | 1.026 | **0.042** | 1.001 | 1.052 | 1.009 | 0.406 | 0.987 | 1.032 |
| Ki67 (% of Tfh) | 1.088 | 0.053 | 0.999 | 1.185 | 1.085 | **0.026** | 1.01 | 1.165 |
| Ki67 (% of Th1-Tfh) | 1.041 | 0.212 | 0.977 | 1.11 | 1.014 | 0.654 | 0.955 | 1.076 |
| Ki67 (% of Th2-Tfh) | 1.078 | 0.077 | 0.992 | 1.172 | 1.091 | **0.012** | 1.019 | 1.167 |

N=207

children. OR: odds ratio; aOR: adjusted OR.

* Adjusted for age as a continuous variable

### Supplementary Material

#### Supplementary Table S5: Relationship between antibodies and odds of symptoms given infection

| | | | Univariate | | | Adjusted | | | |
|---|---|---|---|---|---|---|---|---|---|
| **Target** | **Type** | **OR** | **p** | **95% CI** | | **aOR** | **p** | **95% CI** | |
| MSP2 | IgG1 | 0.5032484 | **<0.001** | 0.39088 | 0.647915 | 1.101556 | 0.394 | 0.881945 | 1.375851 |
| | IgG3 | 0.5032484 | **<0.001** | 0.39088 | 0.647915 | 0.7135221 | **0.02** | 0.536777 | 0.948465 |
| | IgM | 0.609609 | **0.002** | 0.44568 | 0.833835 | 0.8112374 | 0.169 | 0.602301 | 1.092654 |
| | C1q | 0.8766627 | 0.238 | 0.70453 | 1.090855 | 1.08813 | 0.412 | 0.889117 | 1.331687 |
| | FcRIIa | 0.8849889 | 0.264 | 0.71407 | 1.096814 | 1.125994 | 0.254 | 0.918337 | 1.380606 |
| | FcRIII | 0.8355886 | 0.08 | 0.68336 | 1.021721 | 1.071807 | 0.483 | 0.882897 | 1.301137 |
| | OPA | 0.9752543 | **0.039** | 0.95233 | 0.998733 | 0.9965761 | 0.763 | 0.974611 | 1.019036 |
| AMA1 | IgG1 | 0.7731162 | **0.002** | 0.65773 | 0.908746 | 1.000342 | 0.997 | 0.84742 | 1.180858 |
| | IgG3 | 0.6401789 | **0.009** | 0.45722 | 0.896346 | 0.8272131 | 0.235 | 0.604828 | 1.131366 |
| | IgM | 0.3772019 | **<0.001** | 0.23231 | 0.612453 | 0.6424467 | 0.062 | 0.403874 | 1.021946 |
| | C1q | 0.8089297 | 0.089 | 0.63374 | 1.032542 | 1.072248 | 0.542 | 0.857082 | 1.341431 |
| | FcRIIa | 0.7500611 | **<0.001** | 0.64153 | 0.876951 | 0.9508174 | 0.536 | 0.810405 | 1.115558 |
| | FcRIII | 0.7197564 | **0.001** | 0.59268 | 0.874081 | 0.981101 | 0.851 | 0.803646 | 1.1977 |
| | OPA | 0.9782342 | **0.001** | 0.96538 | 0.991257 | 0.9940388 | 0.368 | 0.981186 | 1.00706 |
| CSP | IgG1 | 0.5546815 | **0.008** | 0.35994 | 0.85478 | 0.7542221 | 0.144 | 0.516418 | 1.101533 |
| | IgG3 | 0.8224371 | 0.183 | 0.61678 | 1.096675 | 1.030554 | 0.812 | 0.803895 | 1.32112 |
| | IgM | 0.6637743 | **0.038** | 0.45037 | 0.978302 | 1.037513 | 0.842 | 0.722009 | 1.490887 |
| | OPA | 0.9785899 | **0.001** | 0.96645 | 0.990885 | 0.9911169 | 0.136 | 0.979556 | 1.002815 |
| Pfs230 | IgG1 | 1.293855 | 0.71 | 0.3332 | 5.024239 | 1.602718 | 0.444 | 0.478933 | 5.363385 |
| | IgG3 | 0.2195839 | **0.019** | 0.06157 | 0.783102 | 0.525503 | 0.193 | 0.199387 | 1.385014 |
| | IgM | 0.5397599 | **0.003** | 0.35879 | 0.812006 | 0.9051355 | 0.631 | 0.602777 | 1.359159 |
| | OPA | 0.9665171 | **0.021** | 0.93903 | 0.994805 | 0.9878649 | 0.346 | 0.963083 | 1.013285 |

N=255 children. OR: odds ratio; aOR: adjusted OR.

* Adjusted for age as a continuous variable.

### Supplementary Material

**Supplementary Table S7: Flow cytometry antibodies**

| Target | Clone | Fluorophore | Supplier | Cat number |
|---|---|---|---|---|
| Malaria exposed Tfh phenotyping | | | | |
| CD3 | UCHT1 | AF532 | Invitrogen | 58003842 |
| CD4 | OKT4 | PerCP Cy5.5 | Biolegend | 344635 |
| CXCR5 | J252D4 | BV711 | Biolegend | 356934 |
| PD-1 | EH12.1 | PE-Cy7 | BD Biosciences | 561272 |
| CXCR3 | 1C6 | BV421 | BD Biosciences | 562558 |
| CCR6 | 11A9 | BV650 | BD Biosciences | 563922 |
| CD38 | HIT2 | BV480 | BD Biosciences | 566137 |
| ICOS | C398.4A | APC-Cy7 | Biolegend | 313530 |
| Ki67 | B56 | FITC | BD | 556026 |
| Live Dead | | Zombie NIR | Biolegend | 77182 |
| Malaria naïve Tfh phenotyping | | | | |
| CD3 | SK7 | FITC | Biolegend | 344804 |
| CD4 | OKT4 | PerCP/Cyanine5.5 | Biolegend | 317428 |
| CXCR5 | J252D4 | Brilliant Violet 711 | Biolegend | 356934 |
| PD-1 | EH12.1 | PE-Cy7 | BD Biosciences | 561272 |
| CXCR3 | 1C6 | Brilliant Violet 421 | BD Biosciences | 562558 |
| CCR6 | 11A9 | Brilliant Violet 650 | BD Biosciences | 563922 |
| Live Dead | | Zombie NIR | Biolegend | 423105 |

**Supplementary Material**

**Supplementary Figure S1**

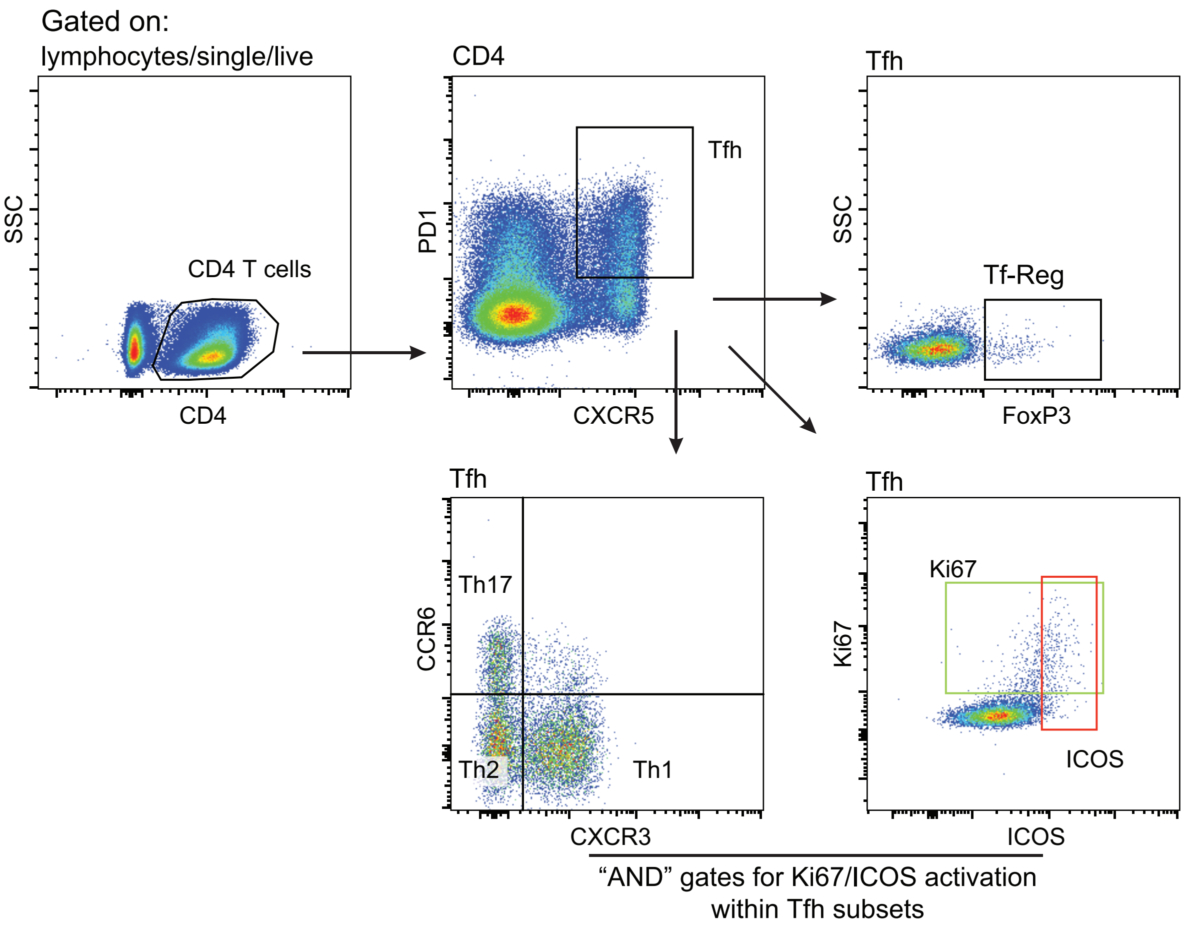

***Supplementary Figure S1: Identification of Tfh***

***A)*** *Gating strategy to identify cTfh, Tf-regulatory (Tf-Reg) cells, subsets and activation. CD4 T cells were gated as CD4+ from lymphocytes/single/live cells. Tfh cells were analysed based on PD1+CXCR5+ cells. Tf-regulatory cells (Tf-Reg) were identified by FoxP3 staining within Tfh cells. cTfh cell subsets were analysed based on CXCR3 and CCR6 staining into Th1 (CXCR3+CCR6-), Th17 (CXC3-CCR6+) and Th2 (CXCR3-CCR6-). Activation was gated as Ki67+ and ICOS+. "AND" gates were made to assess activation markers within subsets.*

### Supplementary Material

#### *Supplementary Figure S2*

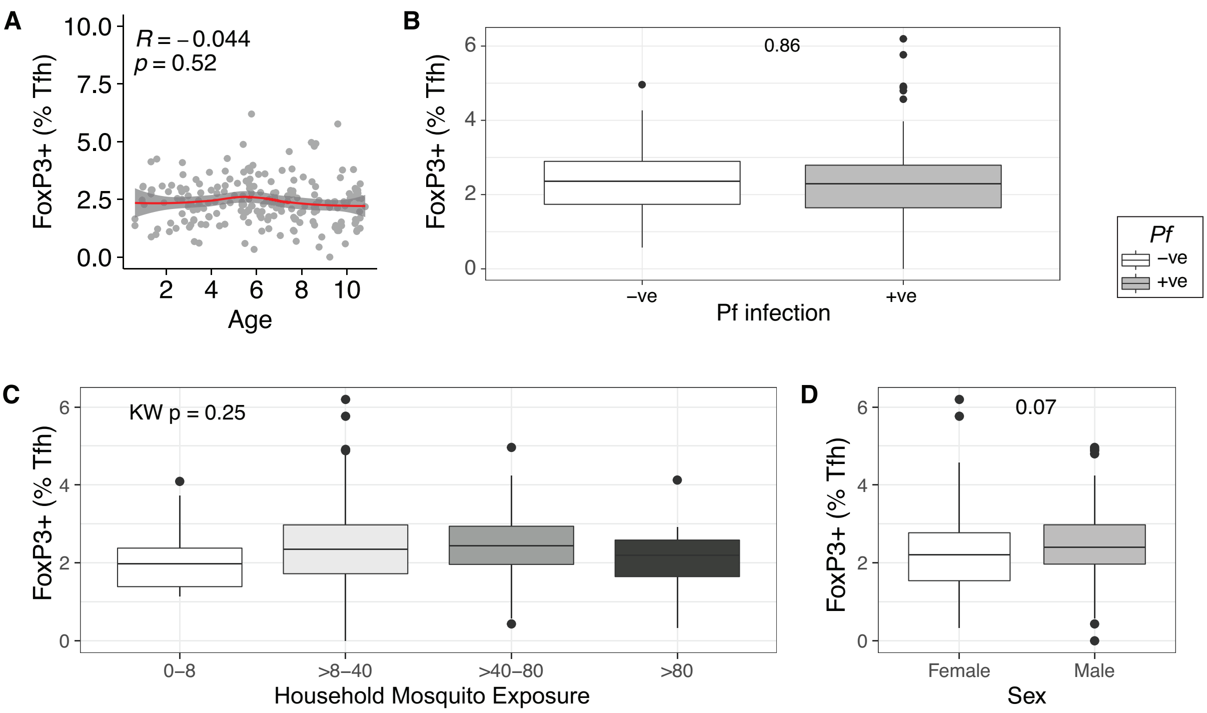

***Supplementary Figure S2: Impact of age and infection on Tf-regulatory cells***
*Tf-regulatory cells were identified as FoxP3+ cells within the Tfh cell compartment.* ***(A)*** *The relationship of the proportion of FoxP3 cells as proportion of Tfh cells with age. Spearmans rho and P indicated.* ***(B)*** *FoxP3 cells and current asymptomatic P. falciparum infection. Mann Whitney U test indicated.* ***(C)*** *Household mosquito exposure (mean mosquitos/house/night). Kruskal Wallis indicated.* ***(D)*** *Sex. Mann Whitney U test indicated.*

### Supplementary Material

#### *Supplementary Figure S3*

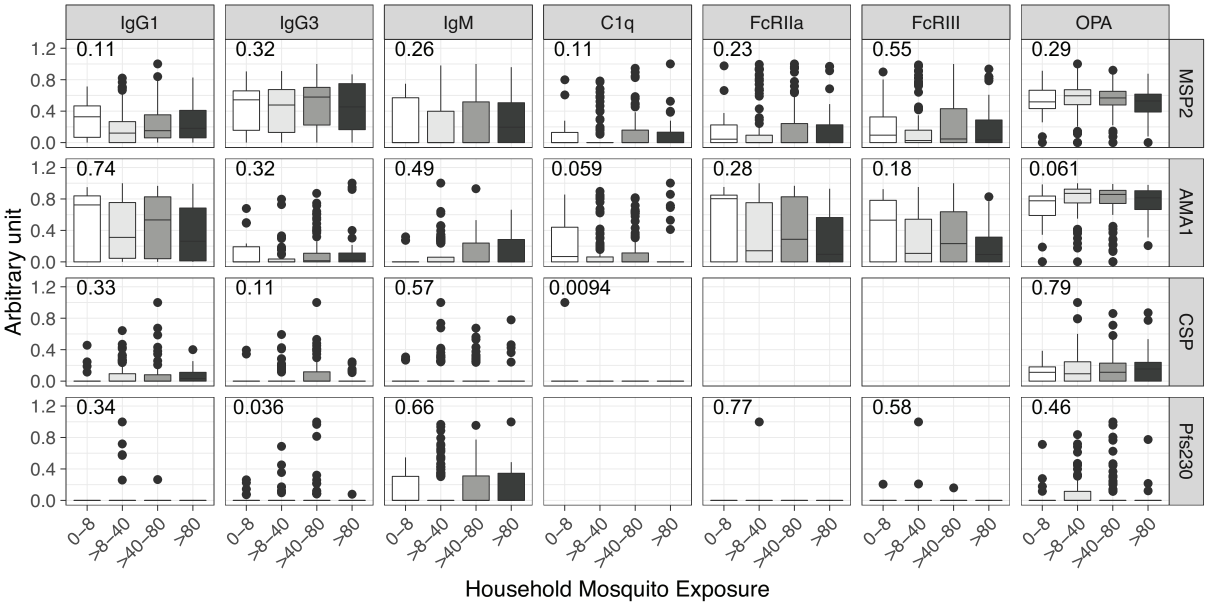

***Supplementary Figure S3: Relationship between antibodies and household mosquito exposure***

*Magnitude of IgG1, IgG3, IgM and functional antibodies that mediated complement fixation (C1q), cross linked FcRIIa and FcRIII and mediated opsonic phagocytosis (OPA) to blood stage (MSP2, AMA1), sporozoite stage (CSP) and gametocyte (Pfs230) antigens with Household Mosquito Exposure (mean mosquitos/household/day). Antibody magnitude is expressed as arbitrary units, which are calculated by thresholding data at positive seroprevalence levels and scaling data to highest responder. Kruskal Wallis P value is indicated.*
